# Easily adaptable head-free training system of macaques for tasks requiring precise measurements of eye position

**DOI:** 10.1101/588566

**Authors:** Katsuhisa Kawaguchi, Paria Pourriahi, Lenka Seillier, Stephane Clery, Hendrikje Nienborg

## Abstract

We describe a modified system for training macaque monkeys without invasive head immobilization on visuomotor tasks requiring the control of eye-movements. The system combines a conventional primate chair, a chair-mounted infrared camera for measuring eye-movements and a custom-made concave reward-delivery spout firmly attached to the chair. The animal was seated head-free inside the chair but the concavity of the spout stabilized its head during task performance. Training on visual fixation and discrimination tasks was successfully performed with this system. Eye-measurements, such as fixation-precision, pupil size as well as micro-saccades were comparable to those obtained using conventional invasive head-fixation methods. The system is inexpensive (∼$40 USD material cost), easy to fabricate in a workshop (technical drawings are included), and readily adjustable between animals without the need to immobilize or sedate them for these adjustments.

**Highlights:**

- We developed an approach to train macaque monkeys head-free on visuomotor tasks requiring measurements of eye position
- The setup is inexpensive, easy to build, and readily adjusted to the animal without the need for sedation
- The system was tested for training on a visual fixation and a visual discrimination task
- Eye measurements (fixation precision, pupil size, microsaccades) were comparable to those from head-fixed animals

## 1. Introduction

Basic systems neuroscience research of cognitive behavior, such as attention, perception and decision-making, often rely on awake macaque monkeys (Roelfsema and Treue, 2014) performing visuomotor tasks. Animals in such studies are frequently trained extensively prior to data acquisition, and many tasks require tight control of the animals’ eye movements (e.g. (Clery et al., 2017)). For precise measurements of eye position the animals’ heads are typically fixed to a primate chair using head-posts surgically implanted to the animal’s skull (e.g. (Adams et al., 2007; Betelak et al., 2001)). To begin the animal’s training using such approaches therefore requires an invasive head-post implantation, and frequently several weeks to months post-surgically for healing and successful osseointegration of the implant into the skull (Betelak et al., 2001) to ensure good stability.

An approach to allow for head-free training without the need for a surgery is therefore desirable not only as a refinement of research with animals from an animal welfare perspective (e.g. (Prescott et al., 2010)), but also because of its potential to accelerate the training procedure, e.g. by taking advantage of the period required for osseointegration for behavioral training. Previous advances with the same goal used molds, helmets or masks covering the animal’s face (Amemori et al., 2015; De Luna et al., 2014; Drucker et al., 2015; Fairhall et al., 2006; Machado and Nelson, 2011; Slater et al., 2016), which were individually tailored to the animals under sedation. Other approaches using transport-boxes of smaller rhesus and new-world monkeys monitored spontaneous gaze direction to natural images in the absence of operant conditioning (Ryan et al., 2019). We focus here on a non-invasive training systems in combination with a primate chair to ease integration into conventional set-ups using head-fixation. We developed a system that is sufficiently flexible that it only requires coarse measurements of the animal’s face, which can be obtained from an animal while seated in a primate chair. The system is integrated in a standard primate chair combined with a commercial eye-tracker. It is inexpensive and simple to build in a standard machine shop (material cost ∼$40USD). We show training data from a visual fixation and discrimination task in one animal as well as detailed measurements of eye movements and pupil size, which were comparable to those obtained from the same animal under head-fixation and two additional head-fixed animals. Because of its flexibility and simplicity the system has the potential to be more widely adapted.

## 2. Methods

### 2.1. Design of the reward spout for head-free training

To train the animal without head-fixation, we used a custom-made concavely shaped reward spout mounted firmly to the chair (schematics and technical drawings in Fig. 1a and c-f, respectively). It consisted of an engineering thermoplastic (copolymere polyoxymethylene, POM-c, “Delrin”, DuPont) milled to a concave cone whose base pointed towards the animal and in whose center a reward tube (OD 4mm, ID 2mm) was inserted and secured with a screw (Fig. 1 c, red arrow). The thermoplastic material was chosen for its sturdiness without being brittle to withstand the animal’s attempts to bite into the rim of the concavity. A wide slit in the downward facing side of the cone ensured that no liquid accumulated inside the concavity of the spout. The blunt top of the cone transitioned to a solid cylindrical part (Fig. 1c). Both conical and cylindrical component were milled in one piece. The depth and diameter of the cone was chosen such that a) it reduced the range of possible head positions from which rewards could be sampled and b) no shadows were cast on the eye to allow for good quality monocular eye-signals. Note that while we designed this spout to approximately match the ventral-dorsal extent (3-4 cm) and width of the upper jaw (4-5 cm) of the animal, our measurements were coarse and did not seem critical. Indeed, we also initially tried a substantially longer spout (87.1 mm instead of 66.9 mm, cf. Fig. 1c), which allowed for overall satisfactory measurements of eye signals although it was more prone to cast shadows on the animal’s eye and hence not used further.

**Fig. 1.**
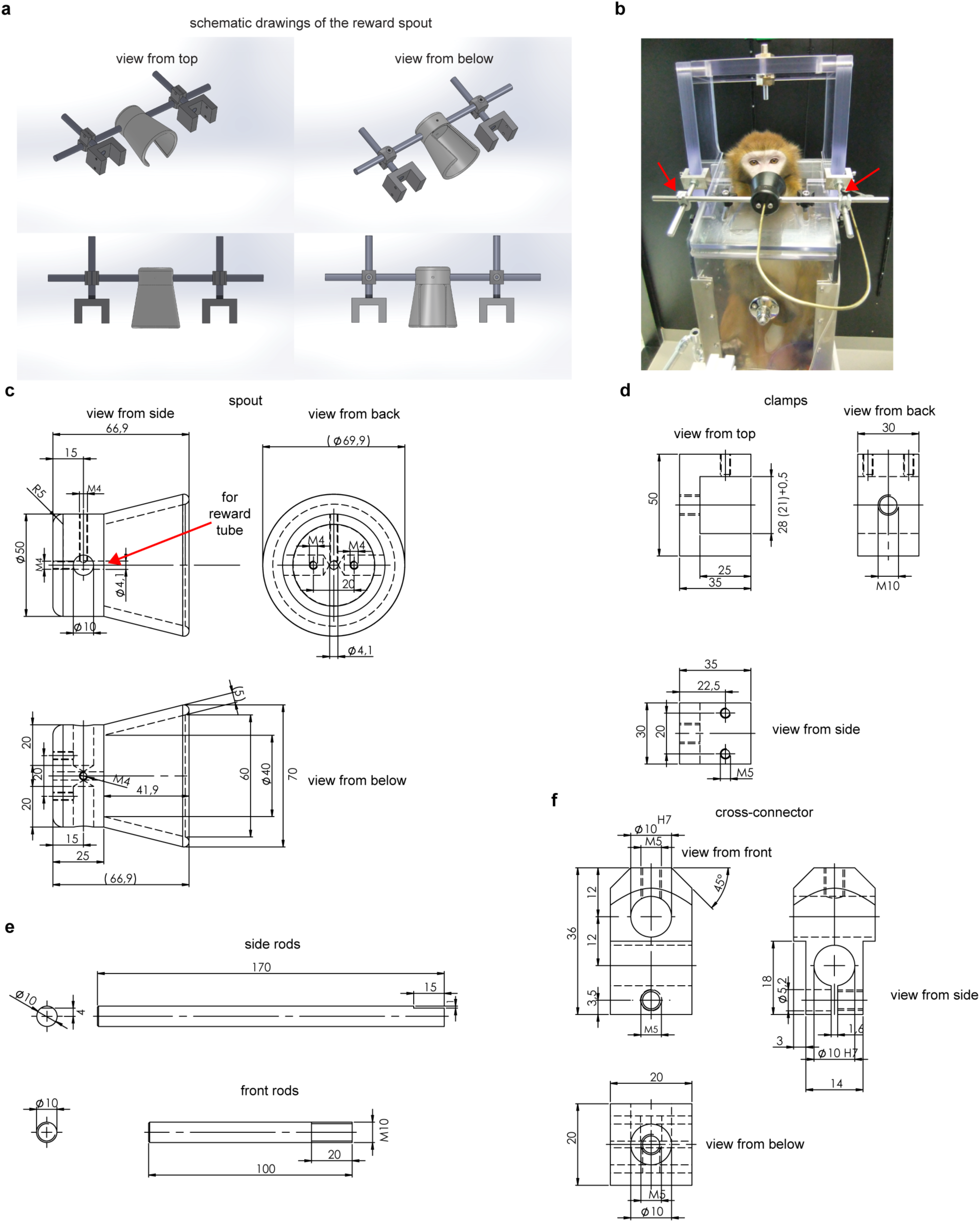
Design of the head-free training system. a) Schematic overview of the spout as well as its mounts to attach it to a conventional primate chair. b) Overview of the system while animal A is being trained. c-f Technical drawings of the components of the system. c) technical drawing of the reward spout milled in one piece out of copolymere polyoxymethylene (cPOM). d) technical drawing of the clamp to mount the spout to the primate chair e) technical drawings of the rods to mount the spout. f) technical drawing of the cross-connectors used to clamp the spout in position.

Moreover, small adjustments could be made by changing the distance by which the reward tube protruded from the bottom of the concavity inside the reward spout. The conical design of the spout in our system was done for convenience of the milling in the manufacturing process. Given the ease to obtain good eye signals with this design there was no need to further refine the shape of the spout. But if, e.g. shadows from the spout led to deteriorated signal quality a narrower shape better tailored to the bridge the noise (resulting in a more triangular cross-section of the spout) could be considered.

The cylindrical part of the spout was screwed to two aluminum rods (OD 10mm), one on each side (Fig. 1e, “front rods”). Note that the spout had to be tightly screwed to the rods to ensure that the monkey could not rotate the spout around the axis of the aluminum rods. These rods (oriented parallel to the front of the primate chair) were then mounted via aluminum cross-connectors (Fig. 1e) to two aluminum rods (OD 10mm, Fig. 1e, “side rods”) that were mounted to the vertical walls of the chair (see Fig. 1b, oriented parallel to the sides of the chair) via aluminum clamps (Fig. 1c). These latter rods remained mounted to the chair between training sessions, while the former and the reward spout were removed between sessions. Only the screws in the cross-connectors (red arrows in Fig. 1b) positioned outside the reach of the animal’s mouth to ensure the experimenter’s safety, therefore needed to be tightened while the animal was seated in the chair. Material cost for all custom-made components was approximately $40 USD.

### 2.2. Measurements of eye-movements and pupil size

The animal’s eye position and pupil size were monocularly measured at 500Hz using an infrared video-based eye tracker (Eyelink 1000, SR Research Ltd, Canada), in the centroid-fitting, pupil-CR (corneal-reflex) and the head-referenced coordinates (HREF) mode. The eye-signals (x position, y position and pupil size) were digitized and stored for the subsequent offline analysis. The eye tracker was mounted in a fixed position on the primate chair (see section 2.3) to minimize variability of the measurements between sessions. For each day’s experiment, we calibrated the eye-tracker by applying a linear transformation (gain and offset) to the raw eye-position signal. This calibration procedure required the animal to fixate at 5 fixation dots (one towards each corner of the screen, 6.6° eccentricity, as well as in the center) sequentially appearing in randomized positions on the monitor. Since the calibration procedure is only possible in animals trained to fixate, we used gain-values obtained from a previously trained animal and manually adjusted the offsets in the initial four fixation training sessions. Our analysis of eye-data focused on the period of animals’ fixation in which the gaze angles were constant.

### 2.3. Camera mount to the chair

To allow for easy daily mounting of the infrared camera in a consistent position without extensive re-calibration we firmly attached an inverted L-shaped mount (aluminum profile) to the primate chair (Crist, Fig. 2b). A custom-made base replaced the commercial base of the camera (“desktop mount model”), and could be easily attached to the horizontal arm of the base. A stopper on the horizontal arm of the base ensured that the camera was mounted in the same position on a daily basis, requiring only minimal refocus to ensure good image quality.

**Fig. 2.**
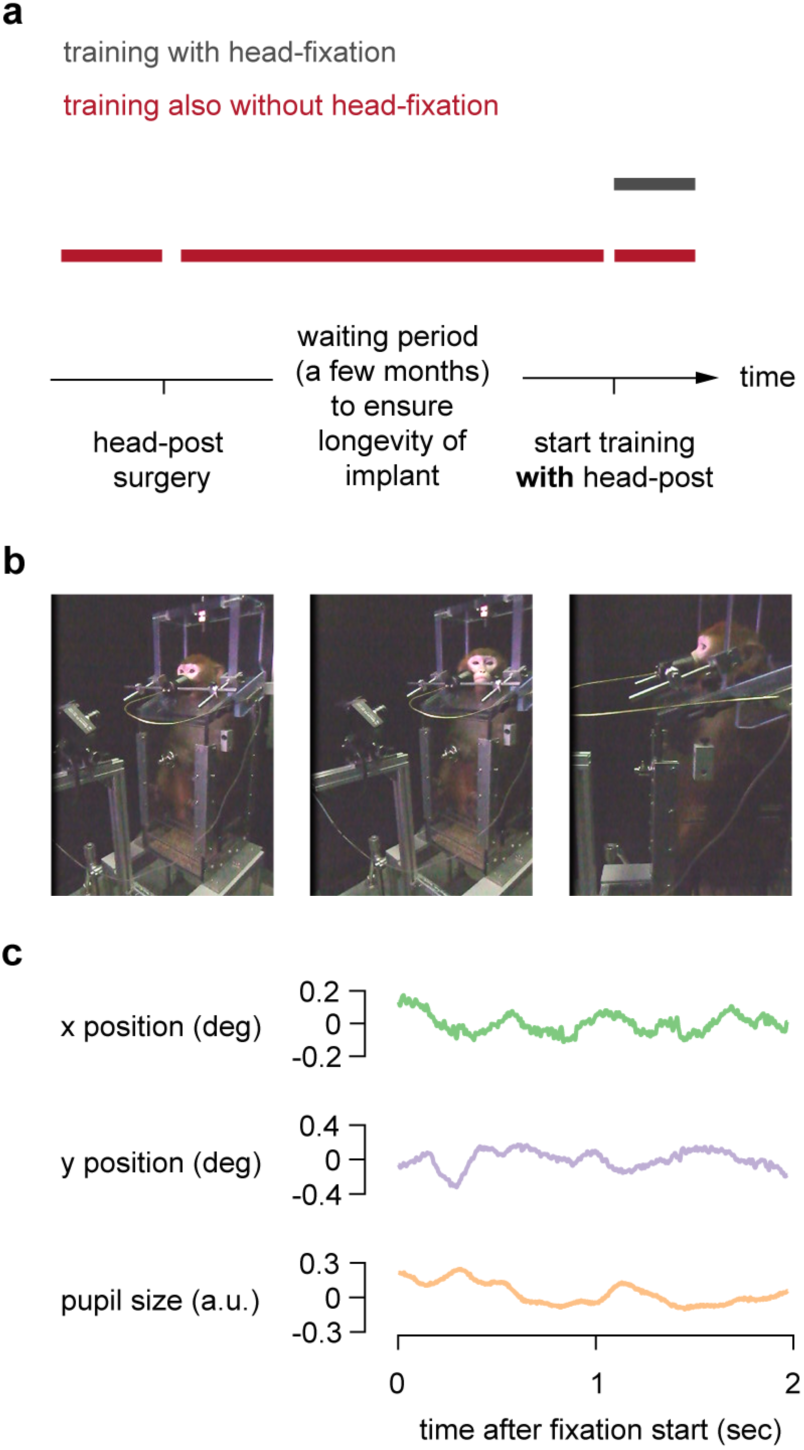
Overview of the approach. a) timeline of head-free training compared to training of head-fixed animals including the period to allow for good osseointegration of the implant to the skull. b) Animal A is being trained using the head-free system. c) Example raw eye-measurements of the x and y-position and the pupil size in one trial.

### 2.4. Animal subjects

This study was approved by the local authorities (Regierungspraesidium Tübingen). We collected data from three male rhesus monkeys, A, M and K (*Macaca mulatta*; K: 6.5kg, M: 8kg and A: 12kg) performing a standard visual fixation task (monkey M and K only under head-fixation and monkey A both head-free and under head-fixation), an orientation discrimination task (monkey A) and a disparity discrimination task (all three monkeys under head-fixation). The monkeys were implanted under general anesthesia with a titanium head-post base on their skull, under their skin, and we developed this head-free training system to take advantage of the post-surgical period for osseointegration in animal A. He was naïve to any behavioral training in a laboratory setting except for climbing into a standard primate chair. After the animal was trained using the head-free system, a metal holder was screwed into the titanium base of the head-holder to allow for conventional head-fixation.

### 2.5. Behavioral training

While monkey M and K received conventional fixation training under head-fixation, monkey A was initially trained using the head-free system. First, he was habituated to the spout of our system and the reward delivery (four sessions). He was then trained on a standard visual fixation task by gradually increasing the fixation duration. We here analyzed the results of all 46 sessions of the fixation training in this animal. Following the fixation training we initiated training on an orientation discrimination task. We initially used the contrast of the distractor target as an additional cue and report here the initial 16 sessions during which both targets were at full contrast such that the only cue the animal could use was the stimulus orientation.

#### 2.5.1 Visual fixation task

The monkeys were required to fixate within a window around the fixation dot 0.1° dva in the center of the monitor to receive juice or water rewards. During the initial training sessions we progressively increased the fixation duration until it reached 2 sec. Once the animals could reliably fixate for 2 sec, we started to present a peripheral visual stimulus (typically a drifting luminance grating) on the screen.

#### 2.5.2. Orientation discrimination task

After animal A learned to maintain stable fixation using the head-free system, we began to train him on a two-alternate forced-choice (2AFC) orientation discrimination task, similar to (Nienborg and Cumming, 2014). Once the animal acquired fixation the stimulus appeared (typically for 2 sec), as well as two choice targets, a horizontally and vertically oriented Gabor, respectively, presented above and below the fixation marker. The vertical position of the choice target was randomized. Once the central fixation marker disappeared the animal was allowed to make his saccade indicating the choice. A saccade to the target whose orientation matched that of the stimulus was rewarded. To discourage the animal from guessing, the available reward size was increased based on his task performance. After three consecutive trials with correct choices, the available reward size was doubled compared to the original reward size. After four consecutive trials with correct choices, the available reward size was again doubled (quadruple compared to the original size) and remained at this size until the next error. After every error trial, the available reward size was reset to the original. For the analyses in Fig. 6b “large available reward” trials refer to both intermediate and large available reward trials collapsed to approximately equalize the number of trials to the small available reward trials.

### 2.4. Visual stimuli

Visual stimuli (luminance linearized) were back-projected on a screen by two projection design projectors (F21 DLP; 60Hz; 1920 × 1080 pixel resolution, 225 cd/m^2^ mean luminance) at a viewing distance of 149 cm (Animal A) or 97.5 (Animal K), or using a DLP LED Propixx projector (ViewPixx; run at 100 Hz 1920!1080 pixel resolution, 30 cd/m2 mean luminance) and an active circular polarizer (Depth Q; 200 Hz) for Animal M (viewing distance 101 cm). Stimuli were generated with custom written software using MATLAB (Mathworks, USA) based on the psychophysics toolbox (Brainard, 1997; Pelli, 1997; Kleiner et al., 2007). For the visual fixation training we used the same set of stimuli as previously described (Seillier et al., 2017), i.e. circular drifting sinusoidal luminance gratings of varying temporal, spatial frequency, contrast and size, randomly interleaved with blank stimuli.

In the orientation discrimination task, the stimuli were 2D Gabor whose orientation and phase was randomly changed on each video-frame (60Hz). Orientation signal strength on each trial was determined according to the probability mass distribution set for the stimulus, analogously to (Nienborg and Cumming, 2009) but in the orientation domain. For the 0% signal stimulus the orientation was drawn from a uniform distribution (8 equally-spaced values between 22.5° and 180°). The monkeys were rewarded randomly on half of the trials on the 0% signal trials. These 0% signal trials were randomly interleaved with horizontal or vertical orientation signal trials. The range of signal strengths was adjusted between sessions to manipulate task difficulty and encourage performance at psychophysical threshold. Typical added signal values were 25%, 50% and 100%.

### 2.5. Analysis

All analyses were performed using Matlab or Python3.

#### 2.5.1 Psychometric threshold

The animal’s choice-behaviors in the orientation discrimination task was summarized as a psychometric function by plotting the probability of ‘vertical’ choices as a function of the signed signal strength *x* and then fitted with a cumulative Gaussian function by maximum likelihood estimation using the fminsearch function in Matlab:

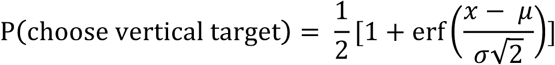

where *erf* denotes the error function, and μ and σ are the mean and standard deviation of the fitted cumulative Gaussian distribution, respectively. The standard deviation σ was defined as the psychophysical threshold and correcponds to the 84% correct level.

#### 2.5.3 Preprocessing of eye traces

We transformed the eye position *x* to velocity *v*, which represented a moving average of velocities over 5 data samples (Engbert and Kliegl, 2003):

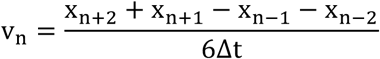

where Δt corresponds to 1/sampling rate. Eye position values were reconstructed using these velocity values to suppress noise (Engbert and Mergenthaler, 2006):

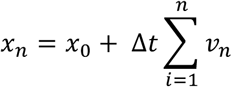

We used the reconstructed eye position for our analyses, where x_0_ is the initial eye position in each session.

#### 2.5.4 Fixation precision

To quantify the fixation quality of the head-free monkey, we computed the variance of the horizontal and vertical eye positions separately. In addition, we computed fixation precision in each session as the fixation span: the area around the mean eye-position during fixation, where the line of sight is found with probability *p* (Cherici et al., 2012). In this analysis we only included trials with successful fixation (typically 2 sec fixation duration). To examine within-trial fixation precision we subtracted the mean eye-position during the fixation period from the eye-position values on each trial. Conversely, to analyze across-trial fixation precision we computed the mean eye position on each trial. We then pooled these eye-position values across all the completed trials in each session and estimated the 2D probability density function by making 2D histograms on a grid covering the entire area of fixation using the Matlab *ndhist* function. Based on this probability density function, we define the area corresponding to the central *75%* of the distribution, as the fixation precision (compare (Cherici et al., 2012)).

#### 2.5.5 Pupil size

Pupil size measurements were z-scored and band-pass filtered as previously described (Kawaguchi et al., 2018). To compare pupil size across sessions, the band-pass filtered pupil size was z-scored using the mean and standard deviation (SD) of the pupil size during the stimulus presentation period across all completed trials within each session. On average the pupil size time-course showed a constriction after the stimulus onset followed by a slow dilation towards the stimulus offset (example in Fig. 6a). To compare the difference in the pupil size between large and small available reward trials, we computed the average pupil size during the 250ms prior to stimulus offset, to compare it to the metric we previously used in head-fixed animals (Kawaguchi et al., 2018). This analysis was restricted to 0% signal trials to exclude potential effects of signal strength on the pupil size analogous to the analysis in (Kawaguchi et al., 2018).

#### 2.5.6 Microsaccades

We used a recently developed microsaccade detection algorithm using a convolutional neural network (https://github.com/berenslab/uneye; (Bellet et al., 2019)). We used the pretrained weights obtained in the original study based on multiple datasets to detect the microsaccades in the head-free animal during the stimulus presentation period. We examined whether the detected microsaccades obeyed the characteristic linear relationship between saccadic peak velocity and amplitude (Zuber et al., 1965), and compared them to microsaccades measured under head-fixation in the same and two additional animals while they performed the analogous task to the orientation discrimination described here in the disparity domain, (see (Kawaguchi et al., 2018; Lueckmann et al., 2018)).

## 3. Results

### 3.1 Training on a standard visual fixation task

We devised this head-free training system to train animals prior to any surgery and to take advantage of the post-surgical period (3-6 months, (Betelak et al., 2001)) aimed at ensuring successful osseointegration of the base-part of a two-part head-fixation implant (Fig. 2a).

We tested this system in one male animal (A) naive to any behavioral training other than to enter the primate chair. The animal was seated in a standard primate chair and trained via operant conditioning to stabilize his head in a fixed position to receive fluid rewards (Fig. 2b, left). Although the animal could move his head freely, the concavity of the reward-spout reduced the variability of the head position whenever he was seeking rewards (Fig. 2b, right) to allow for reliable measurements of monocular eye-position. After four sessions of habituation to the reward spout we were able to start monitoring the animal’s eye position using the chair-mounted video-based eye-tracker during a standard visual fixation task. Example measured eye traces and pupil size are shown in Fig. 2c. Using this head-free system, the monkey was able to fixate successfully for 2 sec within 6 training session, which was somewhat shorter than the number of sessions required in two other animals using conventional head-fixation (13 and 28 sessions in animal M and K, respectively, Fig. 3b). In session seven, following the animal A’s reliable ability to fixate for 2 sec we began to simultaneously present a visual stimulus peripherally during the fixation period. Starting in session eight, we could reduce the size of the fixation window width and height to below 2.5° × 2.5° (Fig. 3c). These increased fixation requirements resulted in a relatively stable proportion of fixation breaks across sessions (Fig. 3d), while the animal worked for increasingly longer sessions over the course of training (Fig. 3e).

**Fig. 3.**
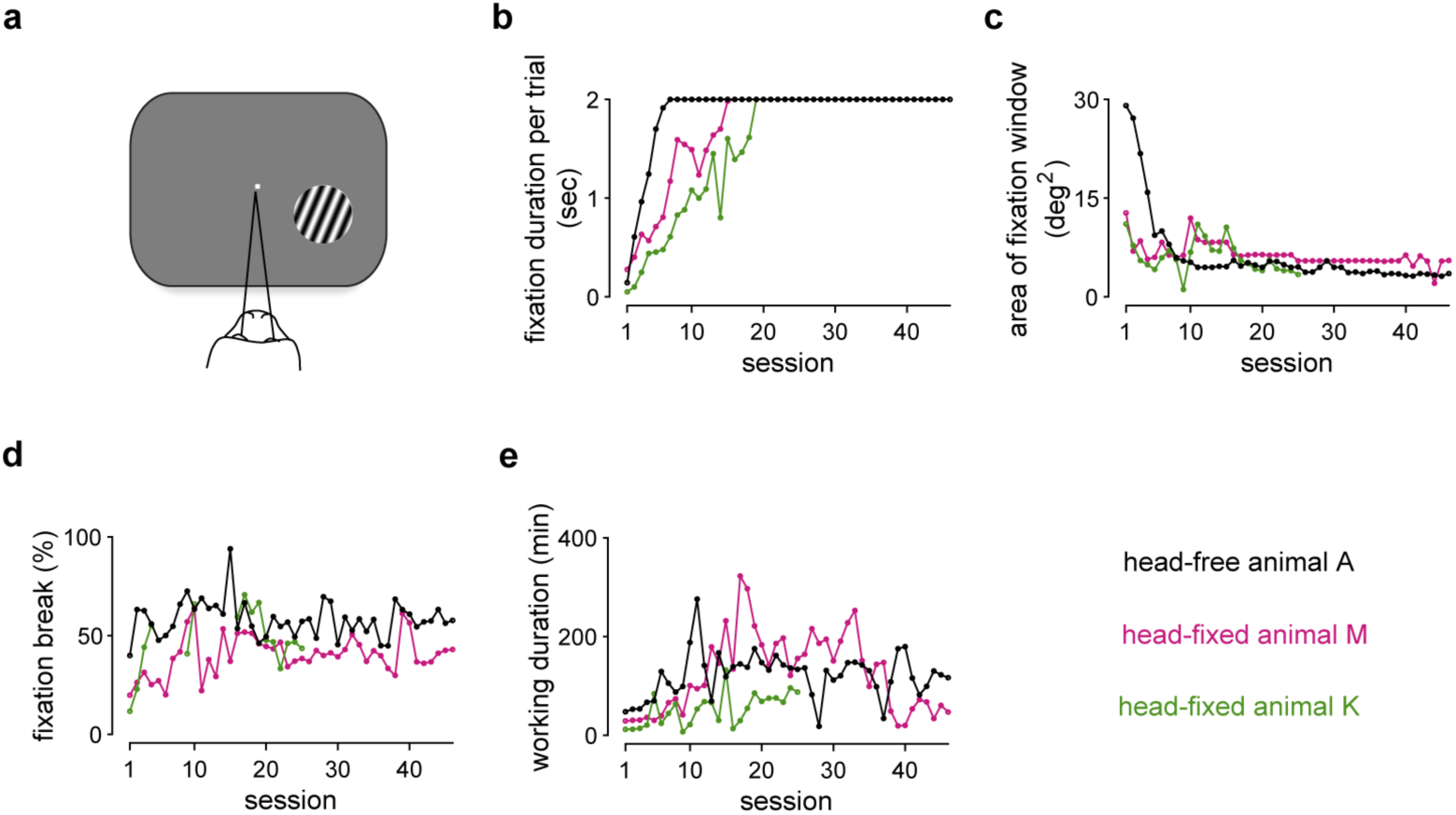
Visual fixation training using the head-free system. a) Schematic of the training setup with the animal fixating on the central fixation marker while a stimulus is presented peripherally. b-e) Black, pink and green data show data for head-free animal A, and head-fixed animal M and K, respectively. The full fixation-training period for head-free training in Animal A are shown, superimposed by partial fixation training in Animal M, and K. Since the training procedure after 25 sessions in animal K deviated from that in animal A and M, we only included the initial 25 sessions of fixation training in this animal. b) The average fixation duration is rapidly increased across consecutive training sessions until the fixation duration of 2 sec is reached. The progression of training of two head-fixed animals is super-imposed. c) The area of the fixation window is progressively decreased across training sessions. d) proportion fixation breaks across sessions was maintained approximately constant. (Note that in animal K eye data are only partially available.) e) The animals worked increasingly longer across sessions.

To examine the quality of the eye-position measurements using this system, we quantified the fixation precision as the variance of eye position (Fig. 4a) and fixation span (Cherici et al., 2012) both within (Fig. 4b) and across (Fig. 4c) trials. We observed that these values improved over the course of training (Spearman’s rank correlation with session number; 4b: r = −0.59, p < 10^-4^ 4c r = −0.76, p < 10^-8^). After about 30 sessions the fixation precision reached values that approached those obtained for fully trained animals under head-fixation (Fig. 4b, c, right). This shows that comparable fixation precision can be reached in our head-free system to that with conventional head-fixation using implanted head-posts.

**Fig. 4.**
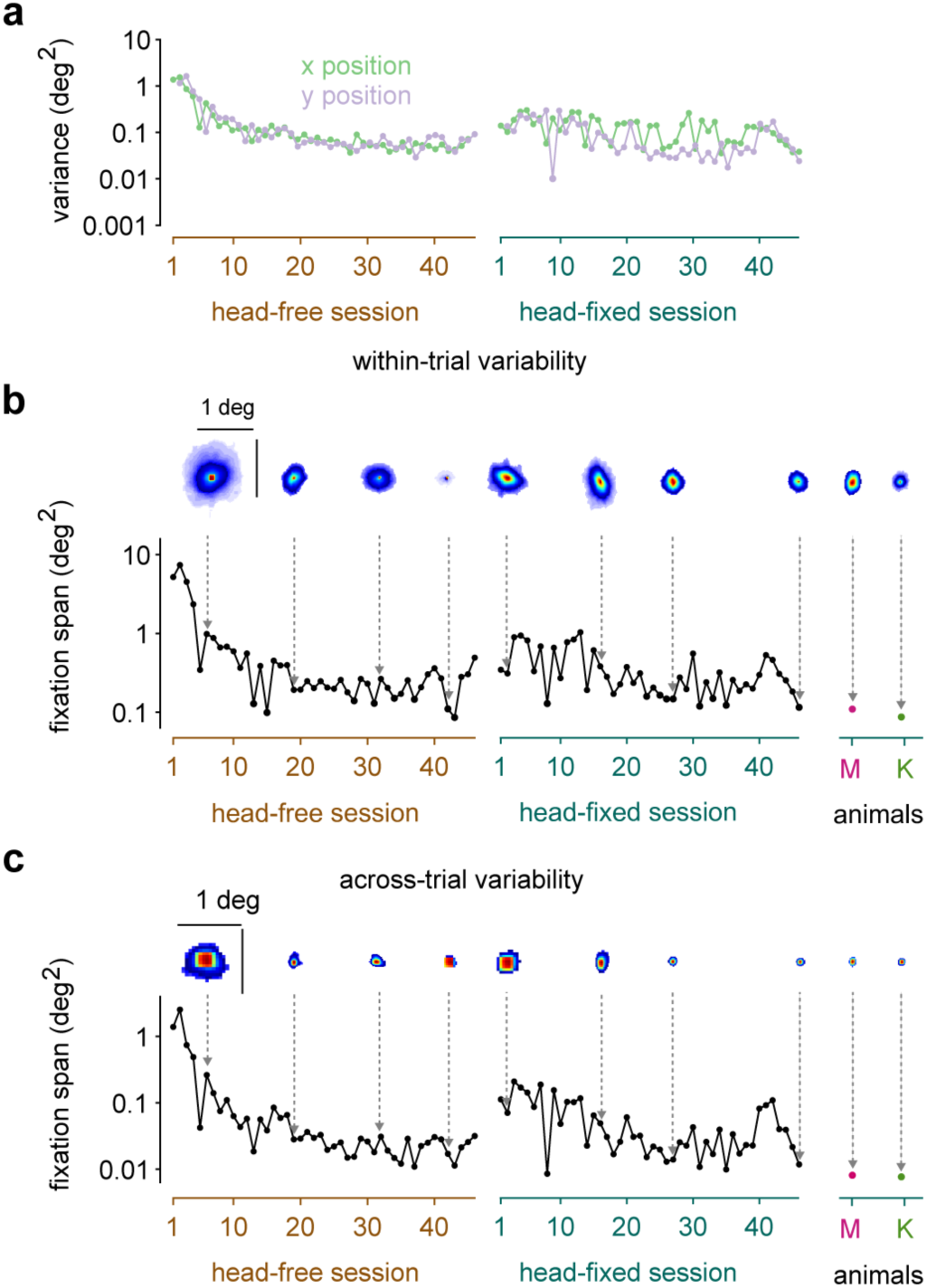
Fixation precision during the fixation task. Left, middle and right column show data for animal A when trained using the head-free system (brown abscissa), animal A when trained under head-fixation (green abscissa), and for fully trained animals M (pink) and K (green) using head-fixation, respectively. a) Variance of x (horizontal; green) and y (vertical; purple) position of the eye. Data points are horizontally jittered for visualization purpose. b) Fixation precision within each trial (b) or across trials (c) quantified as fixation span increased rapidly (smaller fixation spans) with training during the head-free sessions. Insets show the area of the central 75% of the probability density functions (peak normalized for visualization, heat map with red=1) of the gaze positions used to compute the fixation span. Note the transient deterioration in fixation precision after animal A transitions to being head-fixed (middle column). For comparison the fixation precision of two additional fully trained head-fixed animals is shown (right).

### 3.2. Training on an orientation discrimination task

Once the animal achieved good fixation performance we began to train him on an orientation discrimination task (see Methods). During this training we gradually increased the contrast of the incorrect target until the only cue available to the animal to solve the task was the orientation of the stimulus. Here, we analyzed the initial 16 sessions for which both targets were at full contrast such that the animal had to rely on the orientation of the stimulus to solve the task (Fig. 5a). The animal indicated his choice via a saccade to one of the two targets. The video-based eye-tracker captured his choice saccades well (examples from four sessions; Fig. 5b), and the improvement of the psychometric thresholds (Fig.5c) document the training success (Spearman’s rank correlation with session number r = −0.84, p < 10^-4^).

**Fig. 5.**
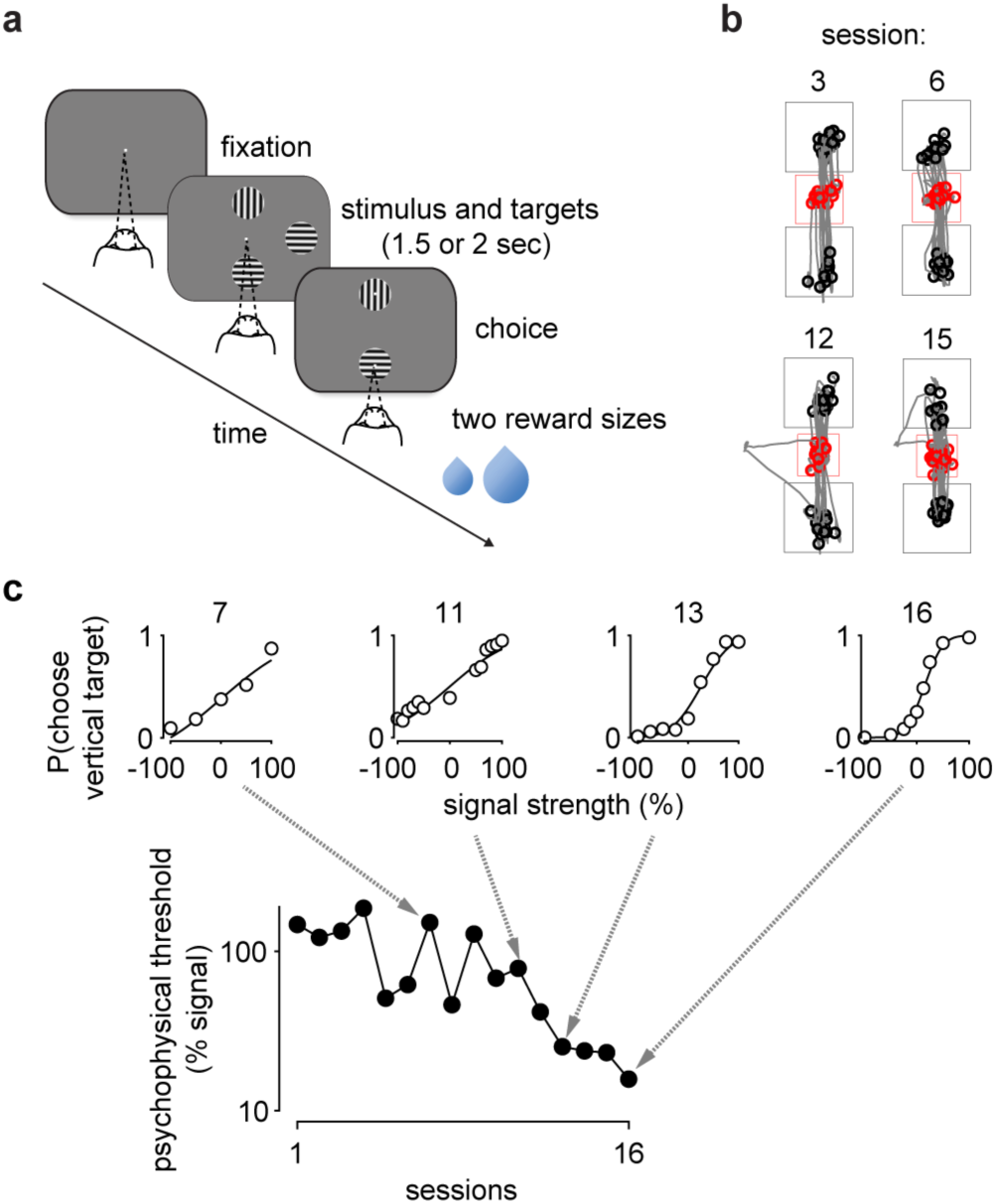
Head-freetrainingonanorientationdiscriminationtask. a) Schematic of the orientation discrimination task. The animal initiated a trial upon fixation. The visual stimulus and the two choice targets were shown for a fixed duration (typically 2 sec). Once the fixation dot was off, the animal made a saccade to one of the targets and received a liquid reward for correct choices. The reward size was changed in a predictable way (see Methods) to discourage guessing. b) Randomly selected choice-saccades from four sessions. Starting points of saccades are in red. End points are in black. Red and gray squares are fixation window and target window, respectively. c) Psychophysical thresholds decreased as a function of sessions, indicating that the psychophysical performance improved. Insets show example psychometric functions from four sessions.

### 3.3 Measurements of modulation in pupil size are comparable to those obtained under head-fixation

We recently used a pupil-size based metric to infer a number of arousal-linked internal states (Ebitz and Platt, 2015; Mitz et al., 2017; Rudebeck et al., 2014; Suzuki et al., 2016), and observed systematic modulation of pupil size with, e.g. available reward size (Kawaguchi et al., 2018). We therefore wondered whether our measurements of the eye signals in the head-free system were of sufficient quality to observe such modulation in this task as well. In the task we used, reward size was changed in a systematic way based on performance (see Methods). We therefore computed the average pupil size in the last 250ms during the stimulus presentation, (as done for head-fixed animals in (Kawaguchi et al., 2018)), and compared this metric for small and large available reward trials across the 16 sessions analyzed here. We found that pupil size was significantly larger in large available reward trials (p = 0.011; Wilcoxon signed rank test; Fig. 6b), very similar to our results in head-fixed animals ((Kawaguchi et al., 2018), their Fig. 3e).

**Fig. 6.**
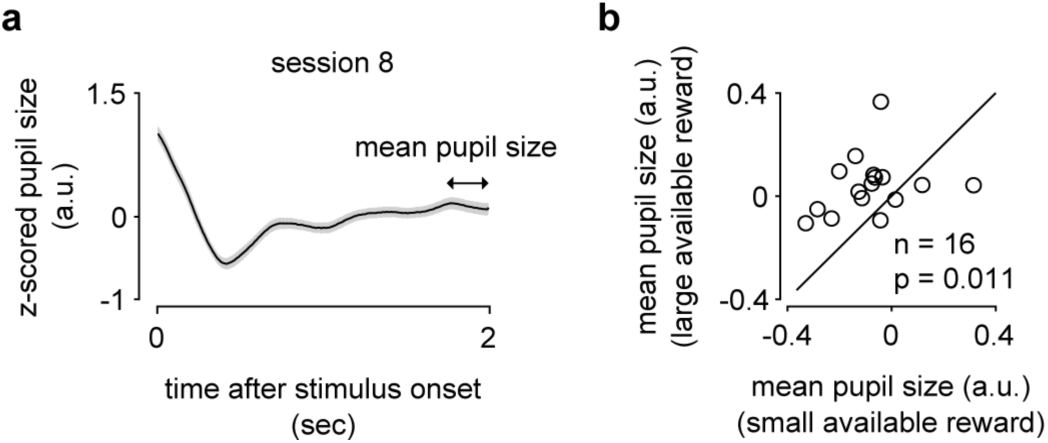
Measurementsofpupilsizemodulationwhenusingthehead-freesystem. a) The pupil size showed a characteristic initial constriction after stimulus onset followed by a dilation towards the end of the stimulus presentation. The average time-course is shown for session 8 (mean ± SEM). b) Modulation in pupil response by available reward size. The average pupil size during the 250ms prior to stimulus offset was significantly larger on large available reward trials was larger (n = 16 sessions, p=0.011; Wilcoxon sign rank test), analogously to our previous findings using the same pupil metric in head-fixed animals (Kawaguchi et al. 2018).

### 3.4 Measurements of microsaccades are comparable to those obtained under head-fixation

Microsaccades are small fixational eye-movements that have been linked to sensory, motor, and cognitive processes (Chen and Hafed, 2013; Herrington et al., 2009; Lowet et al., 2018; McFarland et al., 2015), and are increasingly of interest to cognition research. We therefore wondered whether our eye-position measurements in the head-free system were of sufficient precision to detect microsaccades. The raw eye traces indicate that a recently devised algorithm (Bellet et al., 2019) successfully detected microsaccades using the pretrained weights (see eye traces from two example trials in Fig. 6a). The rate of microsaccades was comparable to that obtained under head-fixation (head-free animal A: 1.14 ± 0.20 s^-1^; head-fixed animal A: 1.64 ± 0.19 s^-1^; head-fixed animal M: 2.19 ± 0.13 s^-1^; head-fixed animal K: 1.23 ± 0.24 s^-1^; mean ± SD) and to values observed in human observers, e.g. (Cherici et al., 2012). Moreover, the microsaccades showed the characteristic linear relationship between peak velocity and amplitude (Zuber et al., 1965), which was similar to that obtained under head-fixation in the same and two additional animals (Pearson correlation: r = 0.61 for 16 sessions, 0.55 for 7 sessions in head-free and head-fixed animal A, respectively; r=0.70 in 8 sessions and r=0.47 in 14 sessions in head-fixed animal M and K, respectively; p<<10^-10^ in all cases, Fig. 7b).

**Fig. 7.**
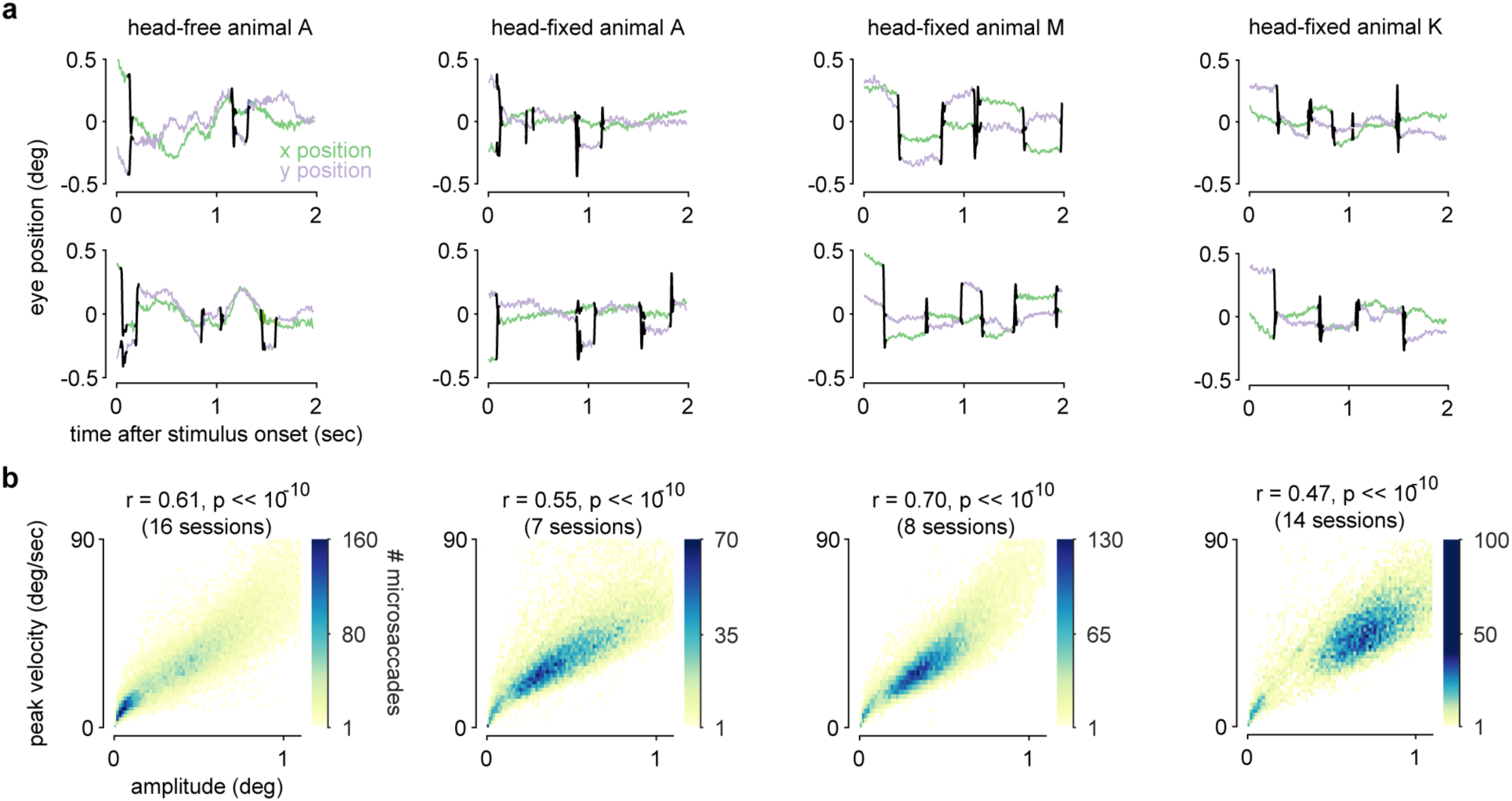
Comparison of microsaccades between the head-free and head-fixed animals. The left two columns show data for animal A for head-free and head-fixed task performance, respectively. The third and forth column show data for head-fixed animals M and K during task performance, respectively. a) Example eye traces with labeled microsaccades (black) for two example trials. b) The characteristic relationship between microsaccade amplitude and peak velocity was also found in animal A in the head-free condition, similar to the head-fixed animals.

### 3.5 Transitioning to head-fixation following the head-free training

After a five-month period of training using the head-free system we transitioned animal A to conventional head-fixation using an implanted head-post. Since the animal had become accustomed to the voluntary engagement in the task and sometimes rotated his body while being seated in the chair we were initially concerned that head-fixation might substantially disrupt his trained visual fixation behavior. But the animal adapted quickly to the head-fixation and required only 10-15 additional sessions of fixation training (Fig. 4b and c, middle, green abscissa) to approach the fixation precision of the fully trained animals that only received fixation training under head-fixation.

Together, these results show that good quality eye-signal measurements can be obtained with this head-free system, allowing for the training on sensorimotor tasks, and that an animal trained head-free can then readily adapt to head fixation.

## 4. Discussion

We described a modified non-invasive system to train macaque monkeys without head-fixation. It has the advantages that, unlike previous non-invasive designs (Amemori et al., 2015; Drucker et al., 2015; Fairhall et al., 2006; Machado and Nelson, 2011; Slater et al., 2016), it does not require sedation of the animals for individual customization and is readily fabricated in a standard workshop. We also performed, for the first time to our knowledge, a detailed analysis of the animal’s eye movements, with a focus on within-and across-trial fixation precision and microsaccades, and pupil size measurements in the head-free system, and compared these to the values observed under head-fixation. We found that the eye-measurements in the head-free animal were overall similar to those obtained under head-fixation in the same animal as well as in two additional animals. The system allows for animal training during the period for osseointegration after implantation of a conventional head-post. The comparison of the time-course of the initial fixation training in one head-free and two head-fixed animals suggests that the head-free training is not slower than conventional training with head-fixation. It may even have the advantage to somewhat accelerate the training process, potentially because it allows for voluntary engagement of the animal in the task, but this would need to be confirmed in a larger sample of animals.

An important observation was that the transition between the head-free system to conventional head-fixation required only minimal re-training, suggesting very little cost in time to employ initial head-free training even when the animal will ultimately engage in experiments requiring head-fixation. This observation, together with the simplicity, flexibility and low material cost should result in a low threshold to adopt this system to more efficiently train the animals as well as improve animal welfare (Prescott et al., 2010), by refining existing set-ups using video-based eye tracking. Finally, the quality of the eye measurements, which was comparable to those obtained under head-fixation, makes the system amenable to combine with neuronal recordings. These may be a tethered configuration with, e.g. chronically or semi-chronically implanted electrodes (e.g. (Ruff et al., 2016)), but would require training the animal to tolerate the touch to the head associated with connecting the cables of the recording system, or wireless recordings (Yin et al., 2014).

## Author contributions

K.K, L.S, P.P, and S.C performed data collection. K.K analyzed data. H.N conceived of the head-free system and supervised the project. K.K and H.N wrote the paper.

## Acknowledgements

This work was supported by a Starting Independent Researcher grant to H.N. from the European Research Council (NEUROOPTOGEN), by funds from the Deutsche Forschungsgemeinschaft awarded to the Centre for Integrative Neuroscience (DFG EXC 307). We are grateful to the animal care staff for all their husbandry expertise and Klaus Vollmer at the workshop in the UKT (Universitätsklinikum Tübingen) for excellent support designing and building the head-free system.

